# Single-molecule conformational dynamics of a transcription factor reveals a continuum of binding modes controlling association and dissociation

**DOI:** 10.1101/2021.03.02.433616

**Authors:** Wei Chen, Wei Lu, Peter G. Wolynes, Elizabeth A. Komives

## Abstract

Binding and unbinding of transcription factors to DNA are kinetically controlled to regulate the transcriptional outcome. Control of the release of the transcription factor NF-κB from DNA is achieved through accelerated dissociation by the inhibitor protein IκBα. Using single-molecule FRET, we observe a continuum of conformations of NF-κB in free and DNA-bound states interconverting on the subseconds to minutes timescale, comparable to *in vivo* binding on the seconds timescale, suggesting that structural dynamics directly control binding kinetics. Much of the DNA-bound NF-κB is partially bound, allowing IκBα invasion to facilitate DNA dissociation. IκBα induces a locked conformation where the DNA-binding domains of NF-κB are too far apart to bind DNA, whereas the loss-of-function IκBα mutant retains the NF-κB conformational ensemble. Overall, our results suggest a novel mechanism with a continuum of binding modes for controlling association and dissociation of transcription factors.

## Introduction

Eukaryotic gene expression starts with a transcription factor binding to a specific DNA sequence. After recognizing the DNA, the transcription factor recruits a vast number of co-activators and eventually RNA polymerase to transcribe DNA to RNA messages (*1*). Transcription occurs as irregular bursts, and the burst frequency and size determine the gene expression level (*2–6*). Live-cell imaging revealed that the burst frequency is controlled by the bound fraction of the transcription factor and the burst size is determined by the transcription factor dwell time (*7*). Bursts continue as long as the transcription factor stays bound and they terminate once the transcription factor dissociates from DNA (*8*). In other words, the rates of association (on rate) and dissociation (off rate) of the transcription factor to and from the DNA determine the amount of RNA produced (*9*). Overall, transcriptional regulation is under kinetic control (*3, 5, 10–12*), where the on and off rates of macromolecular interactions are finely tuned, as opposed to thermodynamic control (*13, 14*), where everything is governed by equilibrium constants.

Recent single-molecule tracking of diffusion and binding of several transcription factors in live cells suggested that the off rate (the inverse of dwell time on DNA) follows a continuous distribution (*15, 16*), instead of only two discrete populations corresponding to specifically and non-specifically bound as implied by decades of biochemical studies (*17*). A continuum of off rates was only found in transcription factors containing intrinsically disorder regions (IDRs), and removal of the IDR resulted in two discrete populations (*16*). It remains unknown whether the broad distribution of off rates reflects the heterogeneous nuclear environment or is an intrinsic kinetic property of the protein-DNA complex (*15, 16*). Yet the essential role of the IDRs suggests that the continuous distribution of off rates may originate from structural heterogeneity in the bound complex.

Controlling the off rate of a transcription factor may seem improbable if it depends on spontaneous dissociation from DNA, in which a higher off rate means fewer contacts and thus less specificity in the protein-DNA complex. Several DNA-binding proteins have been shown to dissociate more readily in the presence of increasing concentrations of the protein itself (*18–23*). The mechanism of this “facilitated dissociation” involves the formation of a transient ternary complex in which two DNA-binding proteins are each bound to part of the DNA-binding site (*24*). For the transcription factor NF-κB, a third player, the inhibitor protein IκBα was shown to facilitate dissociation of NF-κB from DNA in a process later termed molecular stripping (*10, 25*). In this case, a transient IκBα/NF-κB/DNA ternary complex is formed and the off rate is proportional to the IκBα concentration (*25, 26*). Molecular stripping enables tuning the off rate without sacrificing the binding affinity and offers a clean termination of transcription by fast removal of transcription factors on demand (*27*). Similarly, prothymosin α can remove histone H1 from nucleosomes (*28*) and the negative regulator, CITED2 displaces the HIF-1α C-terminal domain from the TAZ1 domain of CBP/p300 (*29*).

A mechanistic understanding of kinetic control, including molecular stripping and the distribution of *in vivo* off rates, demands a deeper understanding of the protein conformational dynamics. To investigate the structural origin of kinetic control for protein-DNA interactions, we carried out single-molecule FRET (smFRET) experiments to visualize the conformational dynamics of NF-κB in real time. Our results revealed a continuum of conformations of NF-κB in both free and DNA-bound states. These conformations interconverted on a range of timescales from hundreds of milliseconds to minutes, comparable to *in vivo* binding events on the seconds timescale (*30*), and suggest a structural origin for the *in vivo* off rate distribution and a binding mechanism utilizing a continuum of binding modes for DNA association and molecular stripping.

Here NF-κB is referred to as the heterodimer formed by the Rel homology domains of subunits RelA (also known as p65) and p50. Each consists of a dimerization domain (DD) and an N-terminal DNA-binding domain (NTD) connected by a 10-amino acid linker. NF-κB recognizes DNA with its two NTDs (**Fig. 1A**), which can undergo nanoscale interdomain motions to expose or occlude the DNA-binding cavity leading to different on rates as shown by hydrogen-deuterium exchange experiments, molecule dynamics (MD) simulation, and stopped-flow fluorescence experiments (*31*). NF-κB activates transcription in response to extracellular stimuli, and is otherwise held inactive in the cytoplasm by the inhibitor protein IκBα (*32*) (**Fig 1B**). Upon stimulation, targeted degradation of IκBα (*33*) allows free NF-κB to translocate to the nucleus and bind regulatory DNA motifs (*34*). NF-κB activates hundreds of genes, including the one of its own inhibitor IκBα, therefore creating a negative feedback loop (*35–37*): the newly synthesized IκBα enters the nucleus (*38*), strips NF-κB from DNA (*10, 25–27*), prevents DNA rebinding, and returns the NF-κB-IκBα complex to the cytoplasm to terminate the transcriptional response (*27, 38*) (**Fig. 1C**). Molecular stripping of NF-κB starts with IκBα encountering DNA-bound NF-κB followed by the formation of a transient IκBα/NF-κB/DNA ternary complex (*10, 26*). MD simulations showed that IκBα induces a twist in the NTDs so that DNA is only held by one NTD, and thus lowers the barrier for dissociation (*10*). An IκBα mutant capable of binding to NF-κB but unable to strip leads to slower export of NF-κB from the nucleus in live cells (*27*), indicating that molecular stripping is critical for efficient termination of the transcriptional response.

**Fig. 1.**
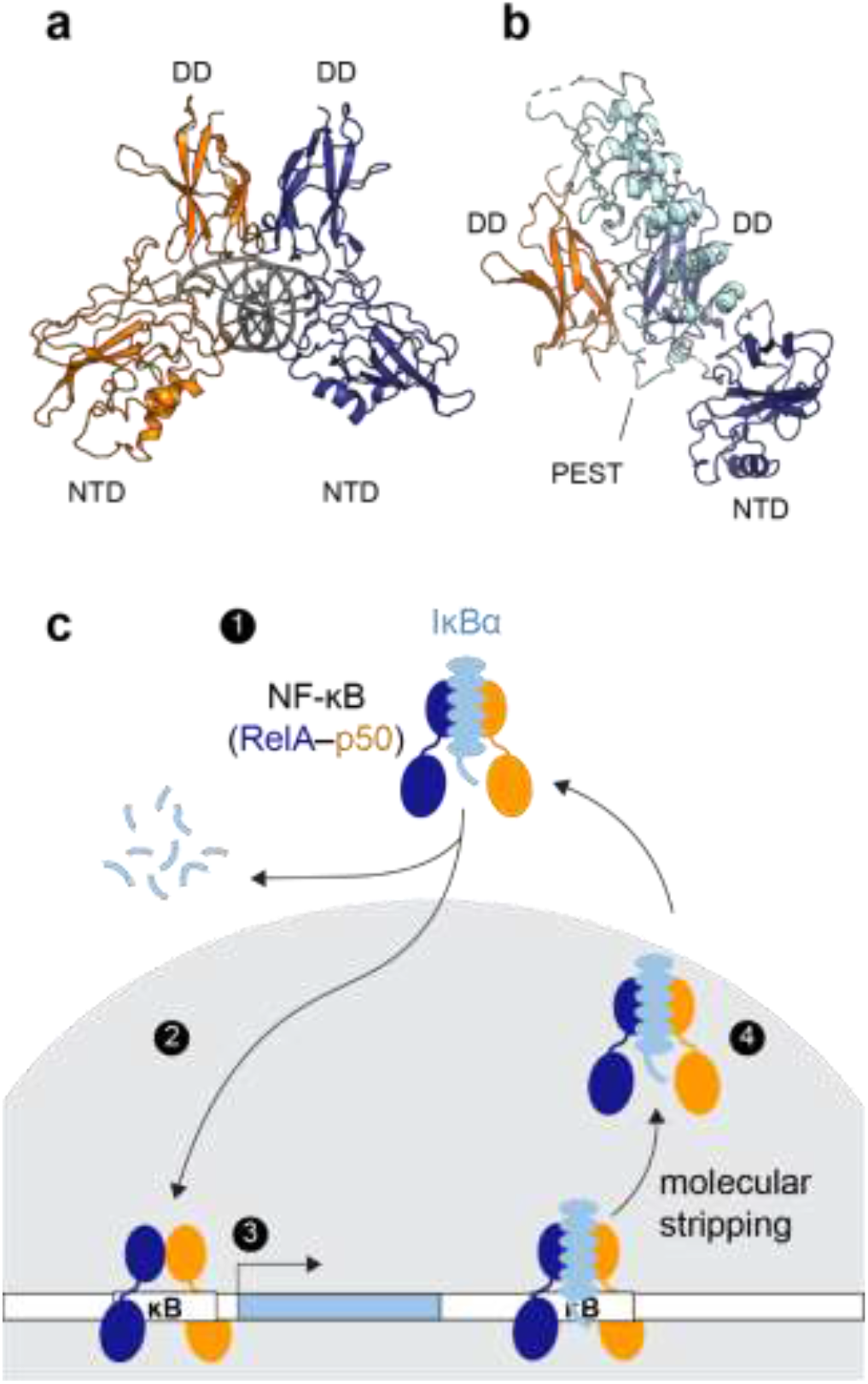
Transcriptional regulation by NF-κB (**A**) Structure of the NF-κB RelA-p50 heterodimer bound to DNA (PDB: 1LE5, RelA in blue, p50 in orange, and DNA in grey). Each monomer contains a dimerization domain (DD) and an N-terminal DNA binding domain (NTD) connected by a 10-amino-acid linker. The NF-κB dimer recognizes its cognate DNA through the two NTDs (**B**) Structure of the NF-κB bound to the inhibitor protein IκBα (PDB: 1IKN, IκBα in cyan). IκBα binds to the DDs of NF-κB. The disordered PEST sequence of IκBα inserts into the DNA-binding cavity. The NTD of p50 was truncated to achieve crystallization. (**C**) Schematic of NF-κB transcriptional regulation showing 1) NF-κB is held inactive in the cytoplasm by IκBα. 2) Upon stimulation, targeted degradation of IκBα allows free NF-κB to translocate to the nucleus (grey). 3) NF-κB binds DNA and activates genes including the one of IκBα, creating newly synthesized IκBα. 4) IκBα enters the nucleus, accelerates NF-κB dissociating from the DNA via molecular stripping, and returns it to the cytoplasm.

Here we report a remarkable observation of a continuum of conformations of NF-κB in the free and DNA-bound states by monitoring the real-time interdomain motions of the NTDs. The broad conformational distribution of DNA-bound NF-κB provides a structural origin of the off rate distribution in live cells (*15, 16*) and a mechanistic understanding of molecular stripping, where the broad distribution containing partially unbound states that allow IκBα to invade the DNA-binding cavity and accelerate DNA dissociation. IκBα induces a single static conformation in NF-κB where the DNA-binding domains are too far apart to both bind DNA, explaining how it accelerates DNA dissociation and inhibits DNA binding (*25–27*). A stripping impaired IκBα mutant still permits DNA binding by allowing the interdomain motions in NF-κB. Overall, our results suggest a novel mechanism of macromolecular interactions by utilizing a continuum of binding modes in both association and dissociation to allow kinetic control. It is also the first time that a continuum of conformations connected by extremely slow motions is directly resolved and visualized, revealing a highly rugged energy landscape and its biological function.

## Results

### Conformational heterogeneity in free NF-κB

Previous coarse-grained (*10*) and all-atom MD simulations (*31*) showed that the two NTDs in an NF-κB dimer can rotate and translate with respect to each other. To experimentally probe the conformational heterogeneity of the NTDs, we carried out TIRF-based smFRET experiments by labeling each NTD with either a donor or acceptor fluorophore. We replaced RelA Gln128 and p50 Phe148 with p-azidophenylalanines (pAzF) by amber suppression and then labeled these two residues specifically with either Alexa Fluor 555 or Alexa Fluor 647 sDIBO alkyne via click chemistry(*39, 40*). The distances between the labeling positions in the NF-κB-DNA crystal structure (*41*) and in an open conformation obtained from previous all-atom simulations (*31*) are 27 Å and 70 Å respectively, corresponding to 0.1 and 1 in FRET efficiency given the 51 Å Förster distance of the dye pair (**Fig. 2A**). The dual-labeled NF-κB was immobilized on a DT20 passivated slide (*42*) through a series of interactions between the C-terminal His-tag on RelA, biotin-conjugated anti-His antibody, NeutrAvidin, and biotin conjugated bovine serum albumin (BSA) (**Fig. 2B**). We collected 426 smFRET traces for free NF-κB. Each smFRET trace covers between 10 and 300 seconds depending on the photobleaching lifetime of the fluorophores. The FRET histogram showed a broad distribution in FRET efficiency from 0.1 to 1 (**Fig. 2C**), revealing a conformational heterogeneity which corresponds to approximate distances of 30 Å to 70 Å between the two NTDs in free NF-κB. The distances we report are close estimates based on rigorous background and donor leakage corrections. Obtaining accurate distances from smFRET requires alternating two-color laser excitation (*43*) and is not the focus of this work.

**Fig. 2.**
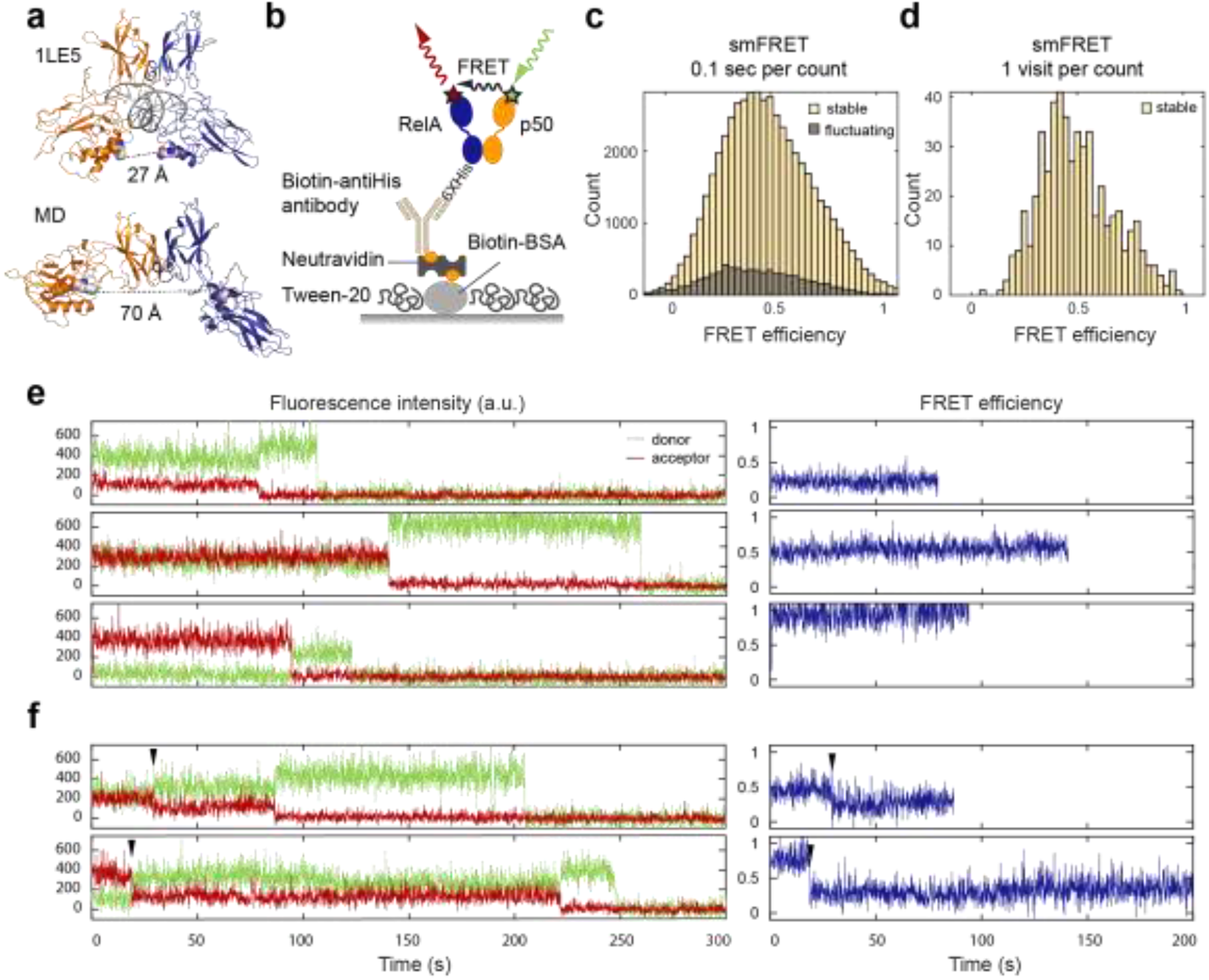
smFRET revealed a continuum of long-lived conformations for free NF-κB. (**A**) In the NF-κB-DNA structure (PDB: 1LE5), the distance between the labeled positions (spheres) would lead to a high FRET efficiency ~1. In MD simulations, free NF-κB could adopt an open conformation, leading to a low FRET efficiency ~0.1. (**B**) Schematic of NF-κB immobilization on a DT20 passivated surface. (**C**) FRET histogram showing a broad distribution of conformations. The histogram was constructed by counting every 0.1 second time window in the first 10 seconds of all traces. (**D**) FRET histogram constructed by counting each visit to a long-lived state suggested a continuum that cannot be separated into a few groups. (**E**) Representative long-lived states with low, mid, and high FRET efficiencies. Sudden drops of the donor or acceptor signals to zero indicate single photobleaching events. (**F**) Transitions (arrowhead) between long-lived states were captured in a subset of traces.

### A continuum of long-lived conformational states in free NF-κB

The smFRET traces showing the time trajectory of single donor and acceptor fluorescence were further classified as stable (82% of the traces) or fluctuating (18% of the traces) (**Fig. 2C**). A stable trace was defined as containing long-lived state(s) with lifetimes longer than the photobleaching lifetime in that trace (**Fig. 2E, F**), which averages 90 sec (**Fig. S1**); a fluctuating trace is characterized by frequent anticorrelation between donor and acceptor signals. In most stable traces, the single NF-κB molecules spent the entire observation time in one long-lived state until photobleaching. Long-lived states with a continuum of FRET efficiencies from low to high were all observed (**Fig. 2E**). To avoid artificial broadening in the histogram caused by noise, in addition to the conventional FRET histogram by binning 0.1 second time windows (**Fig. 2C**), we constructed a histogram for long-lived states by binning the mean FRET efficiency of each visit to a state (**Fig. 2D**). The histogram constructed in this way showed that these long-lived states could not be separated into discrete groups. Instead, we observed a heterogeneous, continuous distribution between high and low FRET efficiencies. Occasionally we observed transitions between two states in a trace (**Fig. 2F**), suggesting long-lived states are interconvertible via slow dynamics on the minutes timescale. A continuum of long-lived states and transitions between them were still observed in the presence of DTT, indicating that the heterogeneity was not caused by the formation of disulfide bonds (**Fig. S3**).

### A continuum of long-lived conformational states in DNA-bound NF-κB

We further investigated the conformational heterogeneity in DNA-bound NF-κB by carrying out smFRET experiments with dual-labeled NF-κB bound to a biotinylated hairpin DNA containing a κB binding site which was immobilized on a passivated surface through biotin-NeutrAvidin interactions (**Fig. 3A**). By immobilizing through biotinylated DNA, we ensured that every NF-κB molecule observed is bound to DNA. We collected 199 smFRET traces for DNA-bound NF-κB, of which 84% could be classified as stable—containing long-lived state(s) with lifetimes longer than the photobleaching lifetime. (**Fig. S1**). Surprisingly, DNA did not shift the conformational ensemble to a single high-FRET state as expected from the crystal structure, but instead shifted the distribution of long-lived states to two major populations including a high-FRET population in good agreement with the structure, as well as an unexpected low-FRET population (**Fig. 3B, C**). Despite these two dominating populations, a continuum of long-lived states including those with mid-FRET efficiencies was observed (**Fig. 3E**,). As was the case for free NF-κB, transitions between long-lived states were captured, suggesting interdomain motions on the minutes timescale (**Fig. 3F**).

**Fig. 3.**
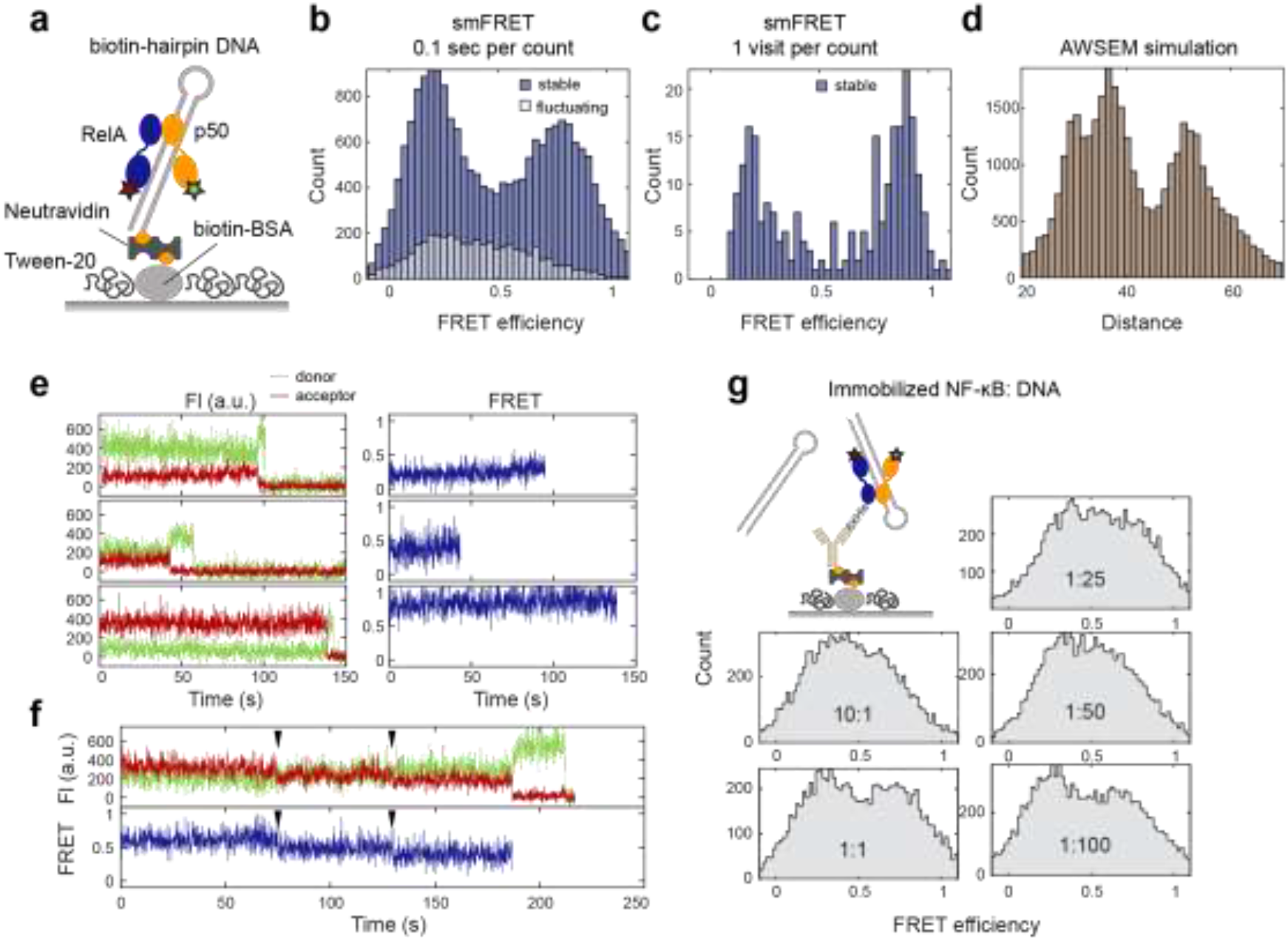
smFRET revealed a continuum of long-lived conformations for DNA-bound NF-κB. (**A**) A biotinylated hairpin DNA with a κB site was used for immobilization through biotin-NeutrAvidin interactions. (**B**) FRET histogram showing two major populations at low and high FRET efficiencies contributed by stable traces. (**C**) FRET histogram constructed by counting each visit to a long-lived state showing a continuous distribution with two dominating populations. (**D**) Distance distribution of between labeling positions by AWSEM simulations showing two dominating populations in DNA-bound NF-κB. (**E**) Representative stable traces showing long-lived states with low, mid, and high FRET efficiencies for DNA-bound NFkB. (**f**) A representative trace showing transitions between long-lived states from high to mid to low FRET efficiencies. (**G**) smFRET titration experiments with immobilized NF-κB and varying DNA concentrations. A series of NF-κB:DNA molar ratios from 10:1 to 1:100 showed similar FRET histograms with a broad band from free NF-κB and two slightly higher peaks at low and high FRET efficiencies from DNA-bound NF-κB.

### Partially bound states within the conformational ensemble of DNA-bound NF-κB

Since all crystal structures of DNA-bound NF-κB indicate a ~30 Å distance between the labeling positions (PDB entries: 1VKX, 1LE5, 1LE9, 1LEI, 2I9T, 2O61, and 3GUT), we were surprised to observe a major population at low FRET efficiency. A crystal structure of NF-κB p50 homodimer bound to two double-stranded RNA molecules (*44*) suggested such low-FRET state could be caused by binding to two copies of DNA on the microscope slide, forcing NTDs to spread apart. To test this possibility, we carried out smFRET titration experiments, this time immobilizing NF-κB and varying DNA concentrations in the solution (**Fig. 3G**). If the low-FRET state was a conformation caused by binding to two DNA molecules, the relative peak height for the low-FRET population would increase as the DNA concentration increased. Because the NF-κB is immobilized, we expect a population of free NF-κB to contribute to the histogram. The results showed that a range of NF-κB:DNA molar ratios from 10:1 to 1:100 lead to almost identical FRET histograms. Each histogram appears to have two slightly higher peaks at high and low FRET efficiencies contributed by DNA-bound NF-κB superimposed on the broad distribution from free NF-κB (**Fig. 3G**). The independence of the FRET histogram on the DNA concentration suggests that the low-FRET state was not a conformation resulting from binding to two copies of DNA but is likely a conformation with one NTD dissociated from DNA.

We turned to coarse-grained AWSEM simulations (*10, 45*) to understand whether NF-κB bound to one copy of DNA could adopt a state with NTDs farther apart leading to low FRET. Indeed, constant temperature simulations revealed a similar broad distribution in distances between labeling positions with a major population of ~30 Å distance as in the crystal structure and another major population with a larger distance that would lead to lower FRET (**Fig. 3D**). Together, the results suggest that within the broad conformational distribution of DNA-bound NF-κB, there are two major populations corresponding to a fully bound state (high FRET) and a partially bound state with the NTDs too far apart to both engage the DNA (low FRET) respectively.

### Interdomain motions of free and DNA-bound NF-κB on the seconds timescale

Besides the stable traces observed for free NF-κB, 17% of the smFRET traces were classified as fluctuating (**Fig. 2C**), based on the fluctuations between different states during the observation time. In some traces, the motions were large enough to connect low- and high-FRET states, corresponding to a 70 Å to 30 Å interdomain distance change (**Fig. 4A**). The motions displayed a high level of heterogeneity and, as was the case for the long-lived states, could not be described by a few discrete groups. In other traces (**Fig. 4B**), the two domains fluctuated around ~0.25 FRET efficiency (distance ~60 Å), and then transitioned to a ~0.8 FRET efficiency (distance ~40 Å) and fluctuated around that value, showing the hierarchical organization of the energy landscape. We also observed that NF-κB molecules transitioned from a fluctuating state to a long-lived state where they could be temporarily trapped (**Fig. 4C**). This again confirmed the long-lived states are not static but are interconvertible with one another and with other shorter-lived conformational states. The fluctuations were subjected to cross-correlation analysis for the donor and acceptor signals. The cross-correlation monotonically decayed as a function of time, suggesting no periodicity for the interdomain motions, and reflecting the stochastic diffusion-like dynamics of single protein domains (**Fig. 4G**). The cross-correlation for fluctuations in free NF-κB could be fitted with a bi-exponential function giving decay rates of 2.0 ± 0.3 s^−1^ and 0.12 ± 0.01 s^−1^ and amplitudes of 22 ± 2 and 38 ± 1 respectively. Characteristic fluctuation times, defined as the inverse of the decay rates, were 0.5 seconds and 8.4 seconds. We also examined the effect of ionic strength on the interdomain motions. A series of smFRET experiments with NaCl concentrations from 100 mM to 250 mM showed that the percentage of fluctuating traces (**Fig. S4A**) and fluctuating time scale (**Fig. S4B**) are dependent on ionic strength but not with a simple monotonic trend.

**Fig. 4.**
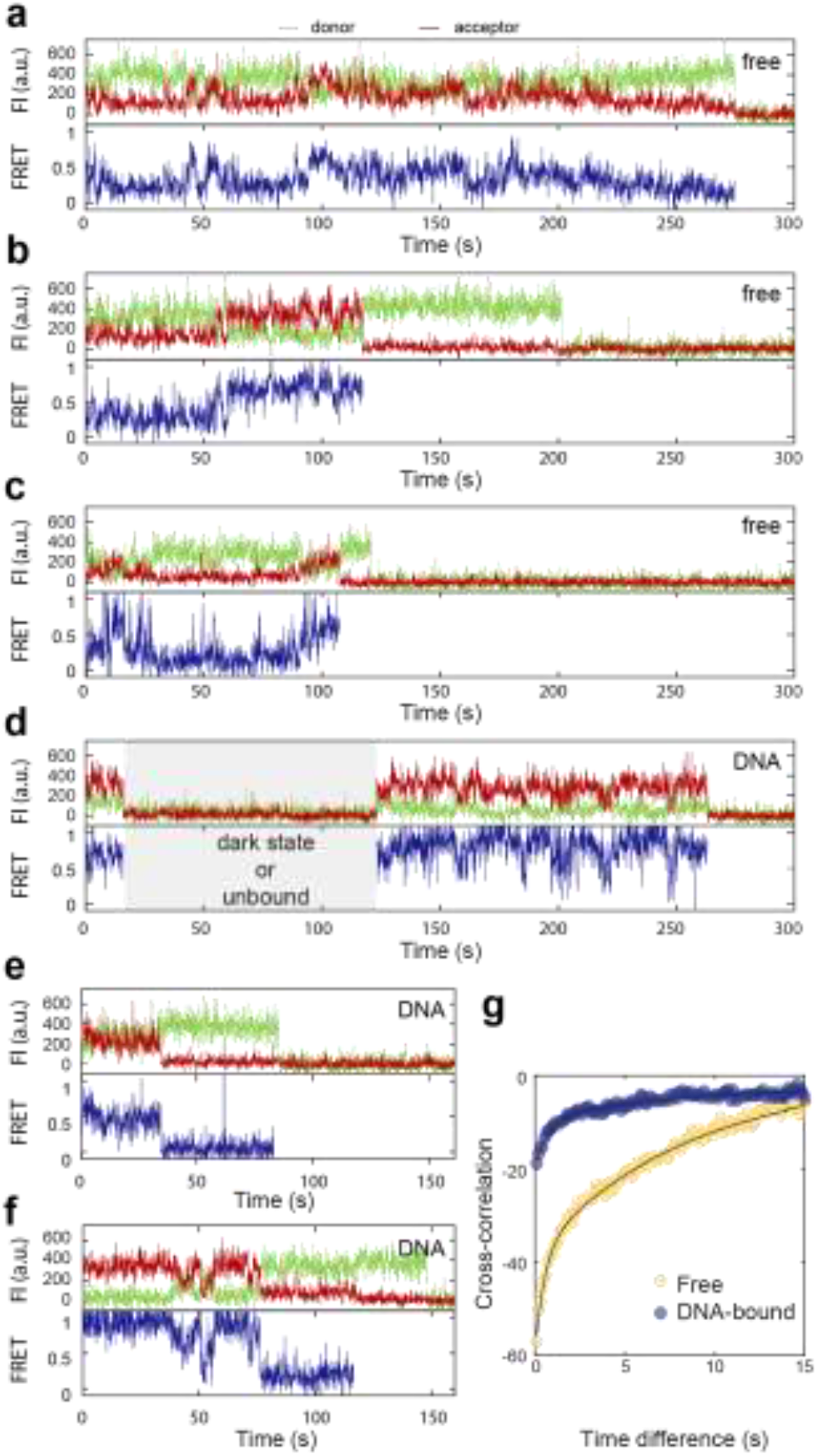
Interdomain motions of free and DNA-bound NF-κB on the seconds timescale. (**A**) A representative fluctuating trace for free NF-κB with fluctuations connecting low- and high-FRET states, which corresponds 30 - 70 Å in the interdomain distance. (**B**) A trace for free NF-κB with small fluctuations around a low-FRET state at first. At 60 seconds, the molecule transitioned to a high-FRET state and fluctuated around it. (**C**) A trace with transitions between fluctuations and long-lived states for free NF-κB, suggesting all the conformational states are interconvertible. (**D** and **E**) Representative fluctuating traces of DNA-bound NF-κB showing small fluctuations. The time period with no signals (grey area) could be caused by a donor dark state, an unbound state, or a state in which an unlabeled NF-κB is bound to the immobilized DNA, and does not affect data interpretation. (**F**) A trace with transitions between fluctuating and long-lived states for DNA-bound NF-κB. (**G**) Cross-correlation analyses for fluctuations of free (yellow) and DNA-bound (blue) NF-κB showing that the interdomain motions of the DNA-bound NF-κB have lower amplitudes and slower rates. Bi-exponential fitting revealed the characteristic fluctuation time to be 0.5 s and 8.4 s for free NF-κB and 0.9 s and 17 s for DNA-bound NF-κB.

NF-κB bound to immobilized DNA also showed fluctuations on the seconds timescale. However, large-scale fluctuations connecting 70 and 30 Å interdomain distances were no longer observed (**Fig. 4D, E**). As in free NF-κB, DNA-bound NF-κB also transitioned between fluctuating states and long-lived states (**Fig. 4F**). Cross-correlation analysis of fluctuations in DNA-bound NF-κB yielded decay rates of 1.1 ± 0.2 s^−1^ and 0.06 ± 0.01 s^−1^ with amplitudes of 11 ± 1 and 8.4 ± 0.6 respectively. The corresponding fluctuation times were 0.9 seconds and 17 seconds. The smaller amplitudes compared to those in free NF-κB reflect the lack of large-scale fluctuations. Both times are two-fold slower than those for free NF-κB. We expect that the decreased fluctuation amplitudes and rates for DNA-bound NF-κB are results of the additional electrostatic interactions introduced by the negatively charged DNA.

### Elimination of conformational heterogeneity by IκBα for molecular stripping

To investigate how the inhibitor protein, IκBα, that strips NF-κB from DNA and inhibits DNA binding affects the interdomain motions of NF-κB, we carried out smFRET experiments with dual-labeled non-tagged NF-κB bound to immobilized IκBα (**Fig. 5a**). By immobilizing through His-tagged IκBα, we ensured that every NF-κB molecule observed is bound to IκBα. The FRET histogram and traces showed that every IκBα-bound NF-κB molecule populated a stable low-FRET state (**Fig. 5B, E**). The histogram built by counting each visit to a long-lived state revealed a narrow conformational distribution in the range of 0.1 to 0.4 FRET efficiency (56 to 74 Å in distance) (**Fig. S5A**). All traces contained long-lived states with lifetime longer than the photobleaching lifetime. No anticorrelation of donor and acceptor signals was observed (**Fig. S2C**), suggesting the interdomain motions displayed in free and DNA-bound NF-κB were completely suppressed by IκBα. Transitions between long-lived states were not observed either. The results show that IκBα eliminated the conformational heterogeneity in NF-κB and locked the interdomain motions in a static state consistent with NTDs being spread apart.

**Fig. 5.**
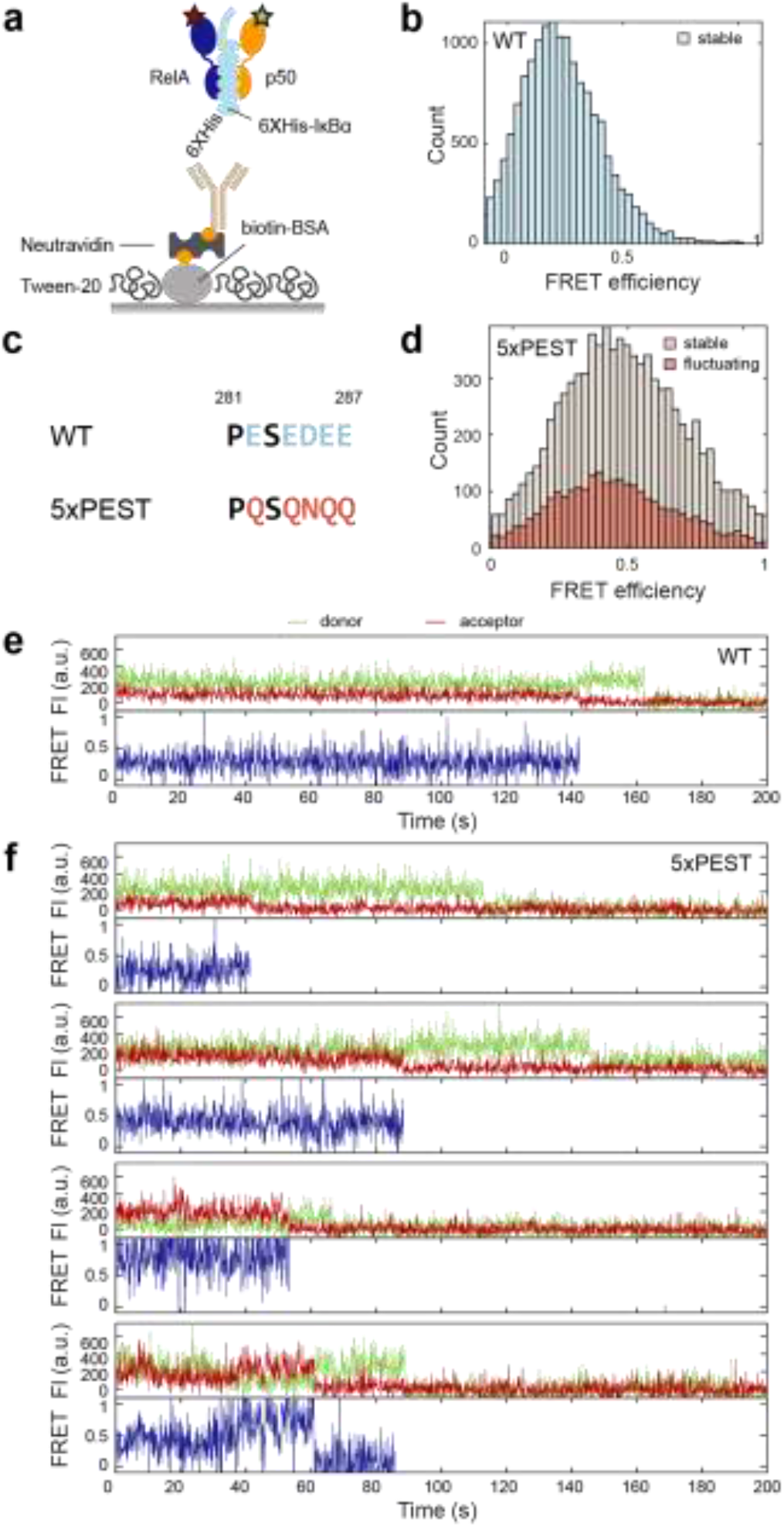
Elimination of conformational heterogeneity of NF-κB by IκBα but not by the 5xPEST IκBα (**A**) The His-tag on IκBα was used to immobilize the NF-κB through the interaction with anti-His antibody biotin conjugate. (**B**) FRET histogram of NF-κB bound to wildtype (WT) IκBα showing only a stable low FRET state was observed. (**C**) The stripping impaired 5xPEST IκBα mutant contains five mutations in the PEST sequence to neutralize negatively charged residues. (**D**) FRET histogram for NF-κB bound to the 5xPEST IκBα showing a broad distribution and a larger proportion of fluctuating traces. (**E**) A representative trace of NF-κB bound to WT IκBα showing the stable low-FRET state. (**F**) Representative traces of NF-κB bound to 5xPEST IκBα showing heterogeneous long-lived states with a range of FRET efficiencies and domain fluctuations.

We next performed the same experiment for NF-κB bound to the 5xPEST IκBα mutant, which contains charge-neutralizing mutations E282Q/E284Q/D285N/E286Q/E287Q in the C-terminal disordered PEST sequence and is not able to strip NF-κB or inhibit DNA binding (*27*) (**Fig. 5C**). The results showed that, in contrast to wildtype, 5xPEST IκBα was not able to lock the NF-κB interdomain motions in a static state. Instead, 5xPEST IκBα-bound NF-κB displayed a continuum of states and domain fluctuations as was observed for free NF-κB (**Fig. 5D, F**). Interestingly, we observed a higher proportion of fluctuating traces in 5xPEST IκBα-bound NF-κB (35%) as compared in free NF-κB (17%), implying that placing a charge-neutral polypeptide in between the NF-κB NTDs reduced the electrostatic interactions that lead to long-lived states. Our results show that the DNA binding function of NF-κB is associated with its interdomain motions, and that the stripping and inhibitory function of IκBα is related to its ability to lock the NF-κB NTDs in a fixed position.

## Discussion

Gene regulation is under kinetic control (*3, 5, 10–12*), in which binding and unbinding of transcription factors to DNA regulate the transcriptional outcome (*9, 15, 16*). To gain a molecular picture of how the on and off rates of transcription factor binding to DNA are controlled, we directly visualized the conformational states of NF-κB and the dynamics of their interconversion at the single-molecule level with TIRF-based smFRET. Our results revealed a remarkable continuum of conformations for NF-κB in the free and DNA-bound states, connected by motions on a range of timescales from hundreds of milliseconds to minutes (**Fig. 6**). The majority of the molecules, either free or DNA-bound, were found to reside in long-lived states interconverting on the minutes timescale, slower than the average *in vivo* dwell time of NF-κB on DNA, which is 4.1 seconds by single-molecule tracking (*30*). Compared to the dwell time of seconds measured *in vivo*, the minute-long conformations of DNA-bound NF-κB would appear static during its residence on DNA, and each of the diverse bound conformations would lead to a unique off rate, resulting in the continuous distribution of rates observed in live cells (*15, 16*).

**Fig. 6.**
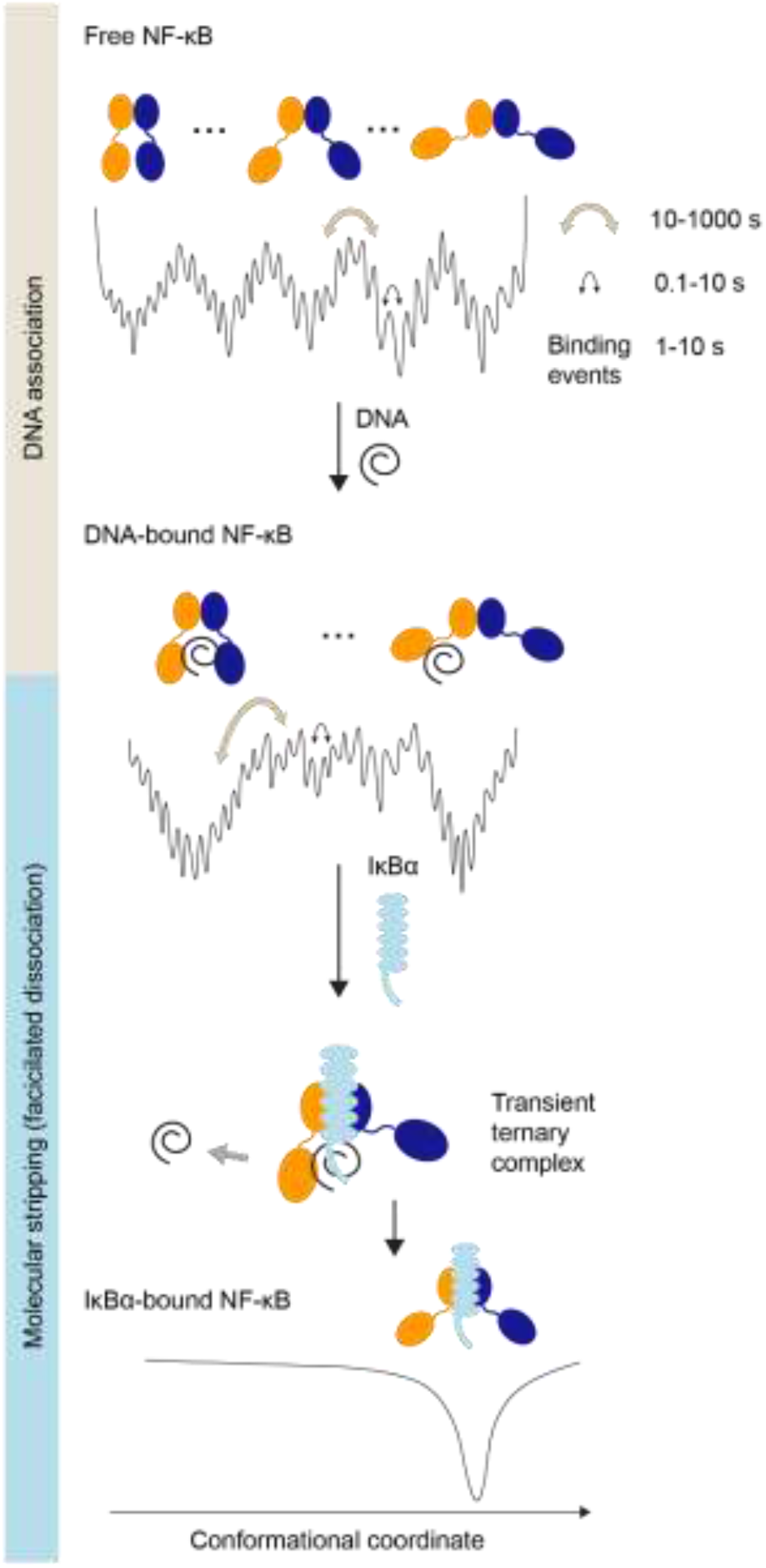
NF-κB utilizes a continuum of binding modes for DNA association and molecular stripping. The energy landscape of free NF-κB is highly rugged and hierarchically organized, resulting in a continuum of conformations interconverting on the subseconds to minutes timescale. The broad distribution of conformations, each with a different degree of DNA accessibility, results in a continuum of on rates for DNA binding. DNA-bound NF-κB also adopts a continuum of distributions, which leads to a continuum of off rates in cells. The two major populations of DNA-bound NF-κB correspond to a fully bound state and a partially bound state; the latter allows the invasion of IκBα to form a ternary complex and eventually strip NF-κB from DNA. Binding of IκBα facilitates DNA dissociation and inhibits DNA binding by inducing a single static conformation in NF-κB where the NTDs are too far apart to both engage DNA.

Similarly, the continuum of conformations in free NF-κB would lead to a distribution of the on rates. The average on rate (k_on_×[DNA]) of NF-κB from single-molecule tracking experiments is 6.7×10^−2^ s^−1^ (*30*), translating to a binding event for an NF-κB molecule every 15 seconds. Again, the long-lived conformations of free NF-κB would appear static from the viewpoint of DNA binding. In previous bulk experiments, we showed that the on rates of NF-κB dimers correlate with the degree of DNA-binding cavity exposure (*31*). The broad distribution of conformations of the NF-κB RelA-p50 heterodimer we resolve with smFRET, each with different DNA accessibility, would result in a continuum of on rates. We also observed more rapid opening and closing of the DNA-binding cavity with fluctuation times of 0.5 and 8.4 seconds for free NF-κB, which would allow for rapid DNA engagement by both domains in the time frame of a single binding event (15 sec).

Our observation of a continuum of conformations sheds light on the mechanism of molecular stripping (*10*), a dissociation mechanism in which the off rate is enhanced by a competitor protein forming a transient ternary complex to lower the dissociation barrier (*10, 26, 27, 46*). Here we showed that much of the DNA-bound NF-κB does not have both NTDs bound to the DNA leaving the DNA-binding cavity vulnerable to the invasion of IκBα. Binding of IκBα eliminated interdomain motions, rigidifying NF-κB into a static state with the NTDs too far apart to both engage DNA. Whereas the 5xPEST IκBα mutant, which is unable to strip NF-κB or inhibit DNA binding (*27*), was unable to elicit the static structure. Our results suggest that the conformational flexibility of NF-κB is required for DNA binding, and that molecular stripping is a process of eliminating such flexibility by IκBα to facilitate the release of DNA and prevent DNA rebinding.

Both conformational flexibility and multivalent interactions together allow facilitated dissociation either by the same protein (*18, 20, 21, 23*) or by a different inhibitor (*25, 28, 29*). These two features enable the visitation of partially bound states and responses to the competitor proteins. Indeed, all the DNA-binding proteins that undergo self-facilitated dissociation are dimers or multidomain proteins with interdomain flexibility and multivalent interactions with DNA (*18, 20, 21, 23*). Likewise, several proteins that undergo inhibitor-facilitated dissociation such as TAZ1/CITED2 and histone H1/prothymosin α are intrinsically disordered with multivalency (*28, 29*). In the case of NF-κB, a continuum of bound states connected by slow interdomain motions allows IκBα to invade the partially exposed DNA-binding cavity and accelerate DNA dissociation. Since nearly all transcription factors are dimers and contain IDRs (*47, 48*), it is likely that many more will display similar continua of conformations and binding rates for exquisite kinetic control.

This is the first time that a continuum of conformations connected by extremely slow motions has been directly resolved and visualized. Historically, navigation through a protein energy landscape has been depicted as interconversion between a few discrete, low-energy states (*49*). Surprisingly, our results reveal a hierarchical rugged energy landscape with many deep wells leading to a continuum of long-lived conformational states interconvertible on the minutes timescale superimposed on a rough surface of substates that interconvert on the hundreds of milliseconds to seconds timescale. Similar findings of multiple continuous conformations and subdiffusive dynamics on a rugged energy landscape have been reported by kinetic analyses of single-molecule experiments and NMR experiments (*50–53*). Yet this is the first time that such slow and heterogeneous motions are visualized in real time. Two features of the NF-κB molecule may be responsible for such slow motions. First, the ionic strength dependence of the slow NF-κB motions suggests they originate from balancing attractive and repulsive interactions from the non-uniform charge distribution. The frustration between the many possible electrostatic interactions would lead to a highly rugged energy landscape with a broad distribution of long-lived conformational states. Indeed, the addition of the negatively charged DNA caused the interdomain motions to be even slower and of reduced amplitude, while the addition of the 5xPEST IκBα mutant with a charge-neutral PEST sequence results in an increased fraction of fluctuating traces. The comparison between wildtype and 5xPEST IκBα also suggests that elimination of the conformational flexibility is achieved through the negatively charged PEST sequence. Second, the highly conserved linker of NF-κB, with the sequence DNRAPNTAEL for RelA and DSKAPNASNL for p50, contains a proline and several charged residues, both of which are known to affect the rigidity and compactness of disordered linkers (*54*) and may lead to slow interdomain motions. In addition, prolines in the linkers can undergo *cis-trans* isomerization on seconds to minutes timescale (*55*) and therefore could also contribute to the slow motions.

In conclusion, our direct visualization of the single-molecule conformational dynamics of the transcription factor NF-κB reveals a continuum of binding modes for DNA association and IκBα facilitated dissociation known as molecular stripping. Our results suggest a novel binding mechanism for kinetically controlling the on rate and off rate of transcription factor binding to DNA. Despite our focus on transcription, this binding mechanism may be prevalent in other aspects of cellular functions where a kinetic control of both association and dissociation is needed.

## Supporting information

supplementary information Chen et al

## Acknowledgments

We thank Hajin Kim at Ulsan National Institute of Science and Technology for his advice on setting up the TIRF microscope, Majid Ghassemian at the Biomolecular and Proteomics Mass Spectrometry Facility of UCSD for carrying out the mass spectrometry analyses, Hannah Baughman for critically reading the manuscript and providing constructive feedback, and Andy McCammon for helpful discussions. W.C. acknowledges fellowship support from the Taiwanese Ministry of Education.

## Data availability

Single-molecule FRET traces are available on https://github.com/komiveslab/NFkB_smFRET_traces.

## Author Contributions

W.C. designed and performed experiments, analyzed data, and wrote the manuscript. W.L. performed and analyzed simulations. P.G.W. supervised simulations. E.A.K. designed experiments and wrote the manuscript.

## Methods

### Protein expression, purification, and labeling

Plasmids of murine RelA_1-325_ (UniProt entry Q04207) and p50_39-363_ (UniProt entry P25799) in pET11a vectors (Plasmid #44744, Addgene) were provided by Gourisankar Ghosh at UCSD. A 6xHis-tag was introduced to the C-terminus of RelA by site-directed mutagenesis for immobilization. Q128 of RelA and F148 of p50 were mutated into Amber stop codons TAG by site-directed mutagenesis for unnatural amino acid incorporation and fluorophore labeling. Plasmid containing RelA or p50 was co-transformed with pEVOL-pAzF (Plasmid #31186, Addgene), the plasmid encoding the aaRS/tRNA pair for incorporating the unnatural amino acid p-azidophenylalanine (pAzF, CAS 33173-53-4, Chem-Impex Int’l. Inc.), into BL21 (DE3) *E. coli* strain (New England Biolabs). Cells were first grown in M9 minimal media with 200 mg/L ampicillin and 34 mg/L chloramphenicol at 37 °C and 180 rpm. At 0.6 - 0.7 OD_600_, the expression of RelA or p50 was induced with 0.2 mM IPTG, and the expression of the aaRS/tRNA was induced with 0.02% L-arabinose. pAzF was added to a 200 mg/L final concentration. Cells were then grown at 18 °C and 180 rpm for 16 hours and harvested by centrifugation at 5,000 rcf for 15 minutes. The cell pellet containing His-tagged RelA was resuspended in 50 mM sodium phosphate pH 8, 150 mM NaCl, 10 mM imidazole, 0.5 mM PMSF with protease inhibitor cocktail (Sigma-Aldrich). Following sonication on ice, the cell lysate was centrifuged at 17,000 rcf for 30 minutes. The supernatant was passed through Ni-NTA resins (Thermo Fisher Scientific) and washed with 10 column volumes of the wash buffer (50 mM sodium phosphate pH 8, 150 mM NaCl, 20 mM imidazole). RelA was eluted with 50 mM sodium phosphate, 150 mM NaCl, 250 mM imidazole.

Elution was pass through a PD 10 desalting column (GE Healthcare) to remove imidazole and eluted with 25 mM Tris pH 7.5, 150 mM NaCl, 0.5 mM EDTA. After added the protease inhibitor cocktail and glycerol (5% final concentration), the sample was aliquoted and stored at −80 °C for further fluorophore labeling and purification. The cell pellet containing non-tagged RelA or p50 was resuspended in 25 mM Tris pH 7.5, 150 mM NaCl, 0.5 mM EDTA, 0.5 mM PMSF. After sonication and centrifugation, the supernatant was passed through SP Sepharose Fast Flow resins (GE Healthcare). An NaCl gradient from 0 to 700 mM NaCl was used to elute the protein. The elution fraction was purified with a PD10 desalting column and stored in 20 mM Tris pH7.5, 150 mM NaCl, 0.5 mM EDTA, 5% glycerol at −80 °C. Reducing agents was avoided to prevent reducing pAzF to pAmF (p-aminophenylalanine). Expression of full-length RelA_1-325_ and p50_39-363_ was confirmed by SDS-PAGE. Site-specific incorporation of pAzF was further confirmed by pepsin digestion and tandem mass spectrometry. Peptides DLE*AISQR (M+H 1093.567, residues 126 – 133 of RelA) and GILHVTKKKV*ETL (M+H 1653.984, residues 138 – 161 of p50) containing pAzF (denoted by asterisk) were identified. Fluorophore labeling was achieved through the copper-free click reaction between the azido sidechain of the incorporated pAzF and the sDIBO functional group on the fluorophore. RelA and p50 were thawed from −80 °C and labeled separately with Click-iT™ Alexa Fluor™ 555 sDIBO Alkyne (FRET donor, Thermo Fisher Scientific) or Click-iT™ Alexa Fluor™ 647 sDIBO Alkyne (FRET acceptor, Thermo Fisher Scientific) with a 1:2 protein-to-dye molar ratio at 4 °C for 2-7 days. The FRET radius R_0_ of the donor/acceptor pair used in this study is 51 Å. Excess dyes were removed with PD10 desalting columns. Labeling efficiency was determined with the extinction coefficients (ε^RelA^_280_ = 20,400 M^−1^ cm^−1^, ε^p50^_280_ = 23,380 M^−1^cm^−1^, ε^Alexa555^_555_ = 155,000 M^−1^cm^−1^, ε^Alexa647^_650_ = 239,000 M^−1^cm^−1^) and the correction factors accounting for A280 from the dyes (CF_Alexa555_ = 0.08, CF_Alexa657_ = 0.03). The labeling efficiency is 29% to 56 % for Alexa Fluor 555 to RelA, 20% to 35% for Alexa Fluor 647 to RelA, 52% to 83% for Alexa Fluor 647 to p50, and 18% to 50% for Alexa Fluor 647 to p50. Labeled RelA and p50 were mixed with a 1:5 donor:acceptor molar ratio to form RelA-p50 heterodimer. The dual-labeled heterodimer was then purified by cation exchange chromatography (Mono S 10/100, GE Healthcare) in 25 mM Tris pH 7.5, 0.5 mM EDTA, 1mM DTT with an NaCl gradient from 100 to 700 mM and size exclusion chromatography (Superdex 200, GE Healthcare) in Tris pH 7.5, 150 mM NaCl, 0.5 mM EDTA, 1mM DTT as previously used for wildtype RelA-p50 heterodimer (*25*).

Human IκBα_67-287_ (Uniprot entry P25963) in pET11a was introduced with a N-terminal 6xHis tag. The 5xPEST IκBα had the following mutations: E282Q/E284Q/D285N/E286Q/E287Q and was expressed and harvested as previously described for nontagged IκBα (*56*). Cell lysates were passed through Ni-NTA resin equilibrated in buffer A (25 mM Tris pH 7.5, 150 mM NaCl, 10 mM imidazole, 10 mM βME), washed with wash buffer (25 mM Tris pH 7.5, 150 mM NaCl, 20 mM imidazole, 10 mM βME), and eluted with buffer B (25 mM Tris pH 7.5, 150 mM NaCl, 250 mM imizadole, 10 mM βME). Fractions containing IκBα were then dialyzed overnight at 4°C in SEC buffer containing 25 mM Tris pH 7.5, 150 mM NaCl, 0.5 mM EDTA, 1mM DTT. The dialyzed protein was then either frozen in 2 mL aliquots at −80°C until needed or immediately further purified from aggregates by size exclusion chromatography (Superdex S75; GE Healthcare) in SEC buffer. The IκBα concentration was determined with the extinction coefficient ε^IκBα^_280_ = 12,950 M^−1^ cm^−1^

### Single-molecule FRET data collection

A prism-type total internal reflection fluorescence (TIRF) microscopy was used. For Alexa Flour 555/ Alexa Flour 647 dye pair, a 532 nm laser (SAPPHIRE 532-300 CW CDRH, Coherent) and a 637 nm laser (OBIS™ 1196625 │ 637nm LX 140mW Laser, Coherent) were set up to excite donor and acceptor respectively. Laser output powers were set to 50 mW for smFRET experiments. 532 nm and 637 nm laser beams were guided by mirrors and a beam splitter to colocalize on the excitation area on the prism. An inverted microscope was set up by mounting the objective (CFI Plan Apochromat VC 60XC WI, Nikon) on a Rapid Automated Modular Microscope System (Applied Scientific Instrumentation). Emission light was further split into donor and acceptor channels with a DV2 Two-Channel, Simultaneous-Imaging System (Photometrics) with band pass filters (ET585/65m for donor channel and ET685/56m for acceptor channel, Chroma). Fluorescent signals were detected by Electron Multiplying Charge-Coupled Device (EMCCD) (iXonEM+ EMCCD camera, DU-897E-CS0-#BV, Andor Technology). DDS-Tween 20 passivated surface (*42*) with 0.2 mg/mL biotinylated bovine serum albumin (A8549, Sigma-Aldrich) and 0.2 mg/mL NeutrAvidin (31000, Thermo Fisher Scientific) were used to immobilize NF-κB. For free NF-κB experiments, NF-κB molecules were immobilized through the 6xHis-tag on the C-terminus of RelA by penta-His antibody biotin conjugate (34440, Qiagen). For DNA-bound NF-κB experiments, DNA-NF-κB complexes were immobilized through the biotinylated hairpin DNA containing the underlined κB sequence:

Biotin-GCATGC<u>GGGAAATTCC</u>ATGCATGCCCCCCATGCAT<u>GGAATTTCCC</u>GCATGC For IκBα-bound NF-κB experiments, IκBα-NF-κB complexes were immobilized through the 6xHis-tag on the N-terminus of IκBα.

Experiments were performed at room temperature in imaging buffer (25 mM Tris pH 7.5, 150 mM NaCl, 0.5 mM EDTA, 0.8 mg/mL glucose oxidase (G2133, Sigma-Aldrich), 0.2 mg/mL catalase (219001, CALBIOCHEM), 0.4% dextrose, and 5 mM Trolox (CAS 53188-07-1, Sigma-Aldrich)). Trolox was prepare by dissolving 10 mg Trolox powder in 10 mL of 25 mM Tris, pH 7.5, 150 mM NaCl, 0.5 mM EDTA with vortexing for 30 minutes and then incubation at room temperature and ambient light overnight. The 637 nm laser was used to confirm the existence of acceptor fluorophores and the 532 nm laser was used for excitation of the donor fluorophore. Data acquisition and analysis software was obtained from the Ha Lab at Johns Hopkins University http://ha.med.jhmi.edu/resources/#1464200861600-0fad9996-bfd4. Single molecule movies were recorded with 100 ms time resolution.

### Single-molecule FRET data analysis

Individual single-molecule traces were extracted from the acquired movies with the IDL scripts in the Ha Lab software package. The traces were further selected based on the criteria: (i) clear single photobleaching step for both donor and acceptor; (ii) anticorrelation pattern for donor and acceptor intensities; (iii) stable total fluorescence intensity before photobleaching; (iii) > 10 sec photobleaching lifetime for both donor and acceptor. Time periods with no signals caused by donor or acceptor dark states, an unbound state, or a state in which an unlabeled NF-κB is bound to the immobilized DNA or IκBα were excluded. For each trace, background was defined as the mean values of the donor and acceptor fluorescence intensities after photobleaching and subtracted from the data. To account for the donor signal leakage to the acceptor channel, we measured the correction factor by collecting data of Alexa Flour 555 labeled RelA. After background subtraction for individual traces of Alexa Flour 555 labeled RelA, the correction factor was defined as the mean value of the ratio of the acceptor intensity to the donor intensity and calculated to be 0.04. Therefore, for a single-molecule trace, the background and donor leakage are corrected by the following equation in which I is the fluorescent intensity:

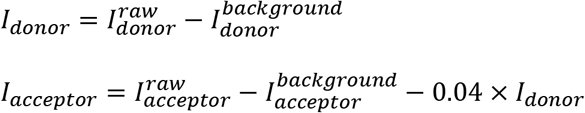

The FRET efficiency (E) was calculated using the corrected fluorescent intensities. E = I_acceptor_ / (I_donor_ + I_acceptor_). Finally, traces were categorized as fluctuating based on whether the signal fluctuations were larger than that caused by noise (shown in the signal after photobleaching). Traces with no fluctuations larger than noise were categorized as stable.

FRET data were presented in two types of histograms. In the first type, the first 1000 timepoints (10 seconds) of each trace were used as data points to build the histogram. In the second type, fluctuating traces were not included, and mean FRET efficiency values of each individual long-lived state were used as data points to build the histogram. The first type of histogram built by counting timepoints is useful for general purposes and the second type built by counting visits to long-live states eliminated noise and was used to show the continuous distribution of states.

Fluctuating traces were subjected to cross-correlation analysis. The cross-correlation function is defined as previously described (*57*). Only traces longer than 30 seconds were included in the analysis and the correlation up 15 seconds were calculated. Results were fitted with a bi-exponential function to compare the characteristic fluctuation time.

### Molecular simulations and analysis

We used OpenAWSEM (*45*) to model the heterodimer-DNA complex. The initial structures of the protein and DNA were from the crystal structure of NFkB (PDB entry: 1LE5). We extended the DNA sequence 100 pairs of A/T at both ends to reduce any finite size effect caused by having a short DNA sequence. The protein and DNA were initially placed far away from each other. Chain A (RelA) of the protein was shifted 300 Å to the left along the x axis from the DNA and chain B (p50) was shifted 300Å to the right. As the simulation began, both chains of the protein were pulled toward the DNA with a native protein-DNA contact bias. This bias was turned off after 10 million steps. Then, we ran another 10 million steps of unbiased simulation. We used a Langevin integrator with a friction of 1 ps^−1^ and a step size of 2 fs. The simulations were carried out at constant temperature of 300 K. The default strength for each term followed our previous study (*58*), with V_contact_=0.75, V_HB_=0.5. We did not use a Gō term in this study. The fragment memory consisted of nine equally weighted structures. Among them, one was the crystal structure, and the other 8 structures are sampled from two independent all-atom simulations (*31*) at 100, 200, 300, and 400 ns. The simulations were repeated 20 times with the same initial structures but different initial velocities and different random seeds. The trajectories were saved every 4000 steps and the last 2000 frames were used for analysis.

