## supplementary information Chen et al for "Single-molecule conformational dynamics of a transcription factor reveals a continuum of binding modes controlling association and dissociation"

**Supplementary Information**
Fig. S1-5

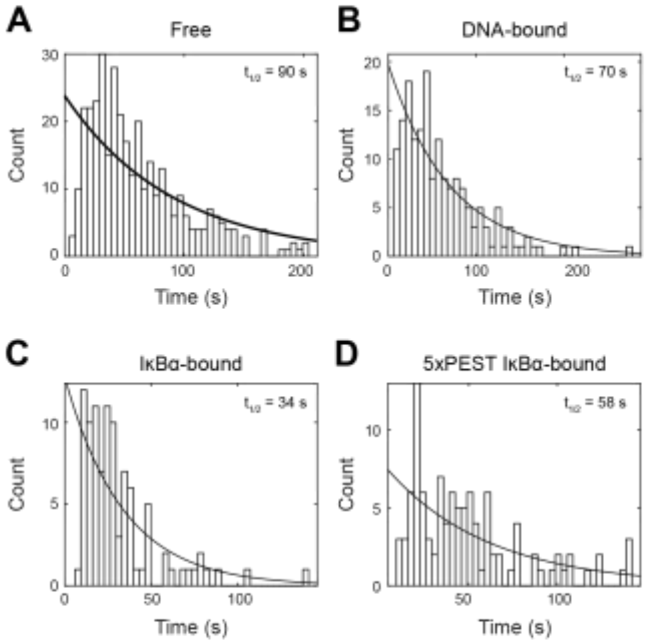


**Fig. S1.** Distribution of photobleaching time for (**A**) free NF-κB, (**B**) DNA-bound NF-κB, (**C**) IκBα-bound NF-κB, and (**D**) 5xPEST IκBα-bound NFkB. The half-life ($t_{1/2}$) of the dye for each set of experiments was obtained from exponential fitting.

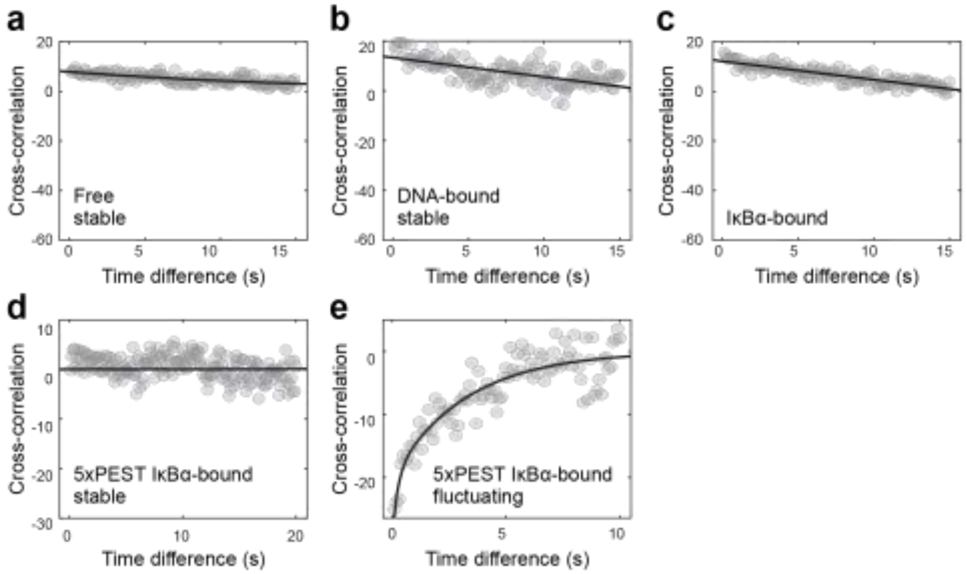


**Fig. S2.** Cross-correlation analyses of smFRET traces. Anti-correlation is indicated by negative values decaying over time. (**A**) No anti-correlation for long-lived states of free NF-κB. (**B**) No anti-correlation for long-lived states of DNA-bound NF-κB. (**C**) No anti-correlation for IκBα-bound NF-κB. (**D**) No anti-correlation for long-lived states of 5xPEST-IκBα-bound NF-κB. (**E**) Anti-correlation for the fluctuating traces of 5xPEST IκBα-bound NF-κB.

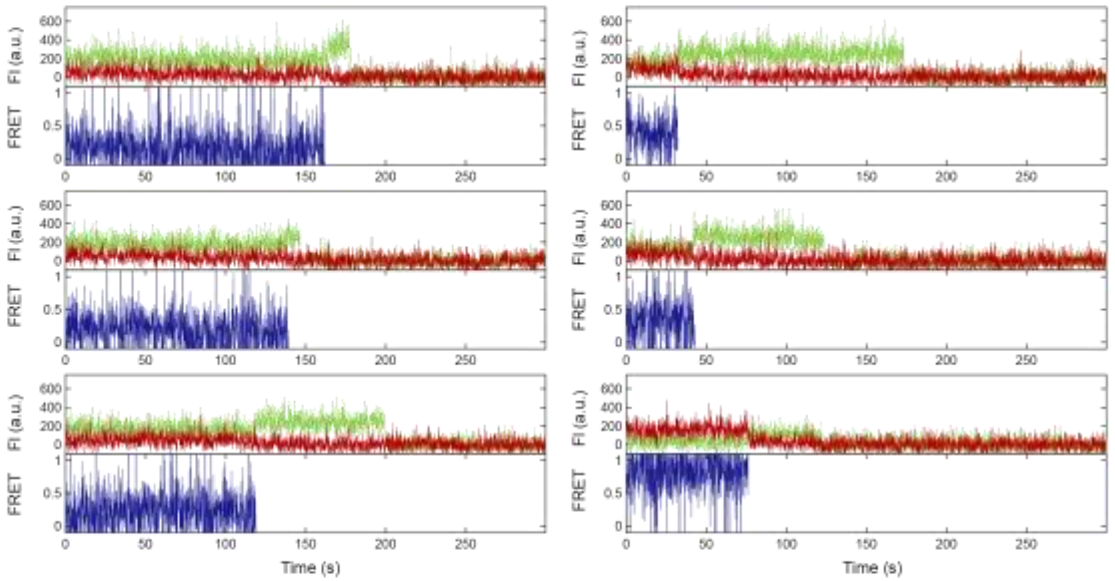


**Fig. S3.** Representative traces for free NF-κB with the addition of DTT. In the presence of DTT, long-lived states with a broad range of FRET efficiencies from low to high were still observed, eliminating the cause of conformational heterogeneity by disulfide bond formation.

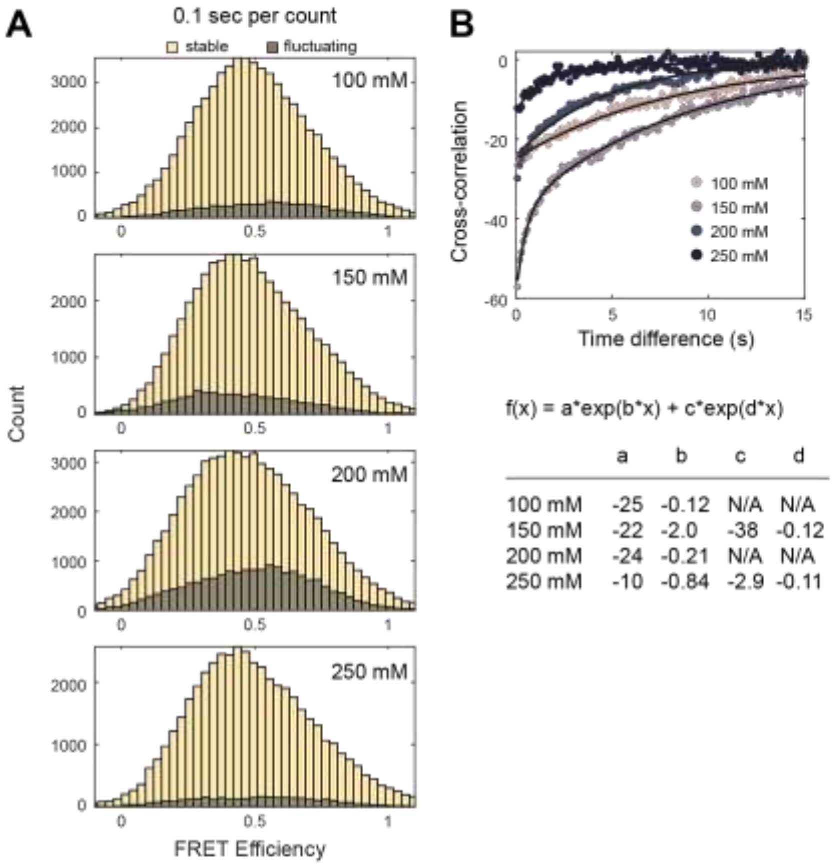


**Fig. S4.** The effect of ionic strength on the conformational dynamics of free NF-κB. (**A**) FRET histograms for free NFκB at different NaCl concentrations from 100 mM to 250 mM. The broad distribution of stable traces was independent of ionic strength. The relative population and the shape of the distribution of fluctuating traces are dependent on ionic strength but not with a simple monotonic trend. (**B**) Cross-correlation analyses on the fluctuating traces and fitting with single or bi-exponential functions showing the fluctuation amplitudes and rates are ionic strength dependent but not with a monotonic trend.

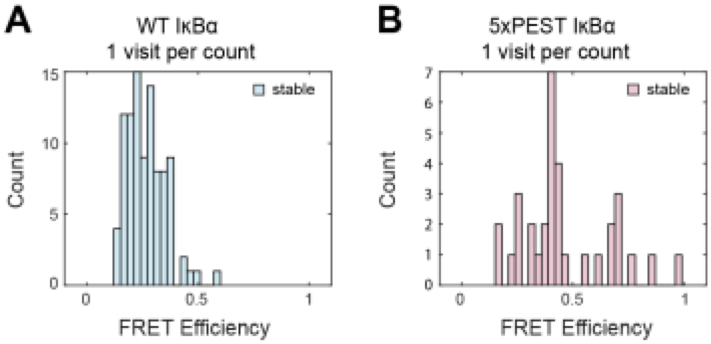


**Fig. S5.** FRET histograms by counting visits to long-lived states for IκBα-bound NFκB. (**A**) NFκB bound to wildtype IκBα adopted a narrow conformational distribution with low-FRET efficiencies. (**B**) NFκB bound to the stripping impaired 5xPEST mutant IκBα adopted a broad distribution of long-lived states.
